# CXCR4 antagonism restores dendritic cell migration and activation in a WHIM syndrome mouse model

**DOI:** 10.64898/2026.05.10.724115

**Authors:** Alexie Ouchakoff, Mathilde Puel, Agnieszka Jaracz-Ros, Molène Docq, Morgan Ocimek, Françoise Mercier-Nomé, Yuna Delarue, Sophie Servain-Viel, Gabriela Cuesta-Margolles, Anvi Laetitia Nguyen, Aurélie Messager, Alain Pruvost, Khadidiatou Kouyate, Katarina Zmajkovicova, Lukas Dillinger, Sandra Zehentmeier, Chi Huu Nguyen, Robert Johnson, Art Taveras, Claire Deback, Patrice Hémon, Françoise Bachelerie, Géraldine Schlecht-Louf

## Abstract

WHIM (warts, hypogammaglobulinemia, infections, and myelokathexis) syndrome is a primary immunodeficiency caused by gain-of-function in CXCR4 chemokine receptor (CXCR4^GOF^) in response to its chemokine ligand CXCL12. The patients suffering from this syndrome display lymphopenia and neutropenia, and most of them show exacerbated susceptibility to human papillomavirus pathogenesis. In a mouse model harboring a WHIM-associated CXCR4 mutation and expressing HPV16 oncoproteins in keratinocytes, we previously reported reduced circulating plasmacytoid dendritic cells (pDCs), mirroring patients’ blood, and impaired dendritic cell (DC) trafficking from the skin to lymphoid organs, with the few migrating DCs displaying an overactivated phenotype. Given the promising results of CXCR4-targeted therapies in WHIM patients, we investigated whether and how the orally available CXCR4-specific antagonist, X4-136, affects DC localization, activation, and trafficking at the subset level, as well as skin immune landscape. CXCR4^GOF^ inhibition corrected defects in circulating myeloid cells and pDCs, as well as in lymph node-resident DCs. Furthermore, it rescued skin DC migration to lymph nodes in WHIM mice, in a context- and subset-dependent manner, by promoting their activation and relocation within the dermis. Taken together, these findings indicate that inhibiting CXCR4^GOF^ may restore skin immunity in WHIM syndrome by rescuing DC counts and functions.

**Key points:**

- CXC R4 gain-of-function inhibition promotes subset-selective dermal dendritic cell migration to lymph nodes in a WHIM syndrome mouse model.
- Inhibiting CXCR4 corrects migratory WHIM dendritic cell hyperactivation with subset-specific effects tied to the inflammatory context.

## Introduction

WHIM syndrome (warts, hypogammaglobulinemia, infections, and myelokathexis) is a primary immunodeficiency most often caused by dominant heterozygous mutations in the carboxyl-terminal region of the CXCR4 chemokine receptor, leading to gain-of-function (GOF) of the CXCL12/CXCR4 axis and impaired receptor desensitization.^1–4^ It manifests as neutropenia combined with lymphopenia and monocytopenia,^5–7^ and most patients develop profuse treatment-refractory human papillomavirus (HPV)-induced warts that can evolve toward carcinoma more often than in the general population.^8–12^

The standard treatments for WHIM patients include granulocyte colony-stimulating factor, supplemental immunoglobulin therapy^11,13^, and prophylactic broad-spectrum antibiotics, but none of these improve HPV-induced lesions^14^. Recent clinical trials have investigated the potential benefits of targeted therapies using two CXCR4-selective antagonists: plerixafor (AMD3100, Mozobil®; Sanofi-Genzyme^15,16^) and the orally available mavorixafor (AMD11070 or AMD-070, X4 Pharmaceuticals^17^). The long-term treatment of WHIM patients by subcutaneous injections of plerixafor twice daily showed encouraging results, including regression of cutaneous warts and HPV-associated carcinomas.^14,18–21^ More recently, phase 2 and 3 studies leading to the FDA-approval of mavorixafor confirmed the beneficial effect of CXCR4 inhibition in WHIM patients, including improvements in lymphopenia and neutropenia, and reduction of HPV-induced cutaneous warts after 18 months of treatment.^22,23^

The immune mechanisms involved in the regression of HPV-induced mucocutaneous lesions remain poorly understood. A contribution of myeloid cells in this process was evidenced in a WHIM patient who spontaneously cured HPV-induced lesions due to a fortuitous deletion of the mutant *CXCR4* allele in one of her myeloid lineage-committed hematopoietic progenitors.^24^ Further data from *FLT3L*-deficient patients suggested that, among myeloid cells, dendritic cells (DCs) would contribute to HPV pathogenesis control,^25^ likely due to their essential role in T-cell activation.^26–29^ Along these lines, we found in a mouse model of WHIM syndrome^30,31^ that CXCR4 GOF (CXCR4^GOF^) limited DC egress from CXCL12-rich environments, including the bone marrow (BM) and dermis.^32^ Thus, plasmacytoid DCs (pDCs) were retained in the BM, accounting for the quantitative defects in circulating pDCs observed in WHIM patients^32,33^, and cutaneous DC migration to skin-draining lymph nodes (SDLNs) was reduced; the migratory DCs (migDCs) that reached these lymphoid organs were highly activated.^32^ This supports the hypothesis that defects in DC functions may contribute to inefficient HPV pathogenesis control in WHIM patients and that rescuing these functions may be instrumental in improving HPV-induced lesions. However, while short-term CXCR4^GOF^ inhibition by plerixafor promoted pDC egress from the BM in a WHIM mouse model, it did not rescue but rather reduced DC migration from skin to SDLNs, whether in the WHIM or control context,^32^ which is consistent with previous observation using another injectable CXCR4 antagonist, motixafortide (4F-benzoyl-TN14003, BL-8040).^34^ These results underscore the role of CXCR4-dependent signaling in DC biology and the need to further investigate the impact of CXCR4^GOF^ inhibition on DCs in preclinical models of WHIM syndrome.

To address this issue, we leveraged a mouse model of WHIM syndrome in the context of HPV-induced chronic skin inflammation to test the effect of 4-week CXCR4 inhibition upon oral administration of the X4-136 antagonist (X4 Pharmaceuticals).^35,36^ The treatment induced elevated numbers of circulating neutrophils, monocytes, DC precursors (pre-DCs) and pDCs in both WHIM and control contexts, as well as an overall rescue of the WHIM-associated DC defects in SDLNs, by increasing pDC, resident (res)DC, migratory type 2 DC (migDC2) and Langerhans cell (migLC) counts. Combining flow cytometry and imaging mass cytometry (IMC), we further showed that this restoration was accompanied by dermal DC2 activation and relocation within the skin of WHIM mice, with limited impact on other cell subsets. Using functional assays, we established that CXCR4^GOF^ inhibition by the X4-136 antagonist promotes dermal DC2 and LC migration to SDLNs, with normalization of their activation levels. This restoration of DC homeostasis suggests that the beneficial effects of CXCR4-specific antagonists on HPV-induced cutaneous lesions in WHIM syndrome patients might, at least in part, result from the rescue of DC distribution and function.

## Methods

### Mice

The CXCR4^+/1013^ (WHIM) mice harboring a heterozygous mutation of the C*xcr4* gene^31^ and the K14-HPV16 (HPV) mice expressing HPV16 early genes under the control of the keratin 14 promoter^37^ (NCI mouse repository) were maintained on the FVB/N genetic background. HPV and WHIM mice were crossed to generate the HPV-WHIM mouse strain^31,32^. Mice were bred under specific pathogen-free conditions in the Animex 2 facility (IPSIT-SFR, UMS-US31-UMS3679). Experiments included 6-week-old males and females, followed French and European guidelines for the use of laboratory animals, and were approved by the local Ethics Committee for Animals (C2EA-26; Animal Care and Use Committee, Villejuif, France).

### Molecules

The X4-136 compound (X4 Pharmaceuticals, USA^35,36^) was prepared in NaCl 0.9% pH 4. X4-136 is an orally bioavailable small-molecule CXCR4 antagonist, and its pharmacological properties are shown in Table 1. Fluorescein isothiocyanate solution (FITC; Merck, France) was prepared at 1 mg/mL in acetone and mixed 1:1 with dibutyl phthalate (Thermo Fischer Scientific, France).

**Table 1.**
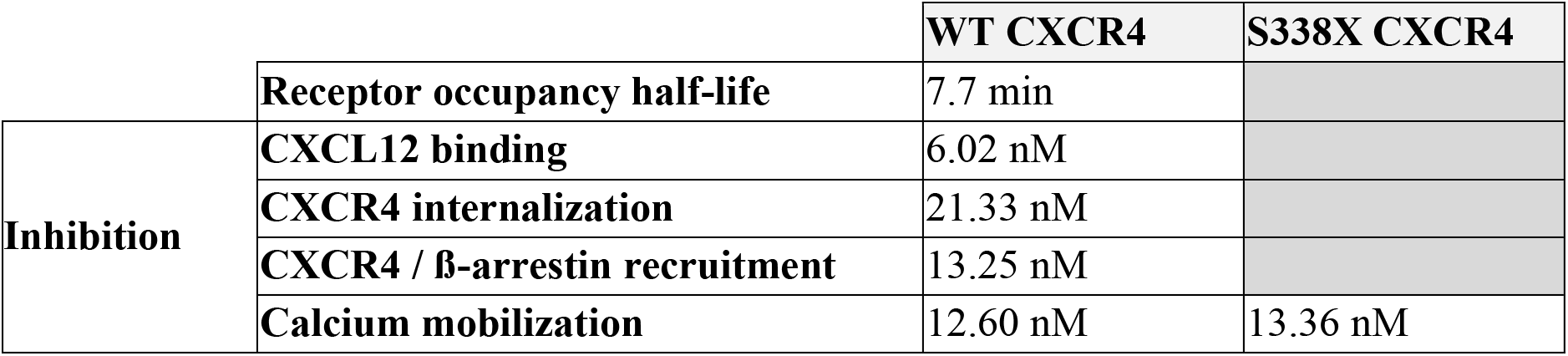
Pharmacological properties of the X4-136 compound.

### Long-term CXCR4 inhibition

HPV-WHIM and HPV mice were treated with the X4-136 compound (0.1 mg/g body weight) by oral gavage 6 days/week for 4 weeks. Control mice received NaCl 0.9% (Sigma-Aldrich, France).

### FITC painting

The flanks of WHIM and control (WT) littermate mice were shaved. The X4-136 compound (0.1 mg/g body weight) was administered per os at time (t)=0. Control mice received NaCl 0.9%. At t=1 h, 200 µL of FITC solution was applied to the mouse flanks. At t=23 h, mice were euthanized, and inguinal LNs were processed for flow cytometry.

### Flow cytometry

Cell suspensions were prepared and analyzed by flow cytometry as described in Gallego et al.^32^ Details are available in the supplemental methods.

### IMC

Data were acquired on a Hyperion imaging system coupled to a Helios Mass Cytometer (Standard BioTools) and analyzed using the Windhager *et al*. workflow^38^. Details are available in the supplemental methods.

### Statistics

Statistical analyses for supervised analysis were performed using Prism 10.2.0 (GraphPad). For data that did not pass the Kolmogorov-Smirnov normality test, the two-tailed Mann-Whitney test or the Kruskal-Wallis test, followed by Dunn’s multiple comparisons test, were used to compare two or several groups, respectively. When normal distribution was validated, Brown-Forsythe and Welch ANOVA tests followed by Dunnett’s T3 multiple comparisons tests were performed. Statistics for unsupervised analysis were performed using R versions 4.5.1 and 4.5.2 in RStudio^39^. Kruskal-Wallis test was performed with Benjamini-Hochberg (BH) correction of p-value.

## Results

### Chronic administration of X4-136 increases blood leukocyte counts in HPV-WHIM and HPV mice

We previously reported that CXCR4^GOF^ causes leukopenia in WHIM mice on the C57Bl/6 genetic background^32^. We first confirmed that HPV-WHIM mice display lower lymphocyte and myeloid leukocyte counts than HPV mice in the FVB/N background (Figure S1). Thus, WHIM-associated lymphopenia and neutropenia were present in this HPV-associated chronic inflammation model. We next tested the effects of chronic CXCR4 inhibition with the X4-136 antagonist upon daily (6 days out of 7) oral administration of the compound for 25 to 27 days (Figure 1A). Blood cell subset counts were analyzed after 8 days of treatment, 22 hours after the final dose, to capture changes in baseline levels. Leukocyte counts were increased in both HPV-WHIM- and HPV-treated mice compared with their respective untreated controls (Figure 1B). Detailed exploration showed no significant difference in lymphoid cell counts between X4-136-treated HPV-WHIM and HPV mice and their respective controls, except for B cells in HPV mice (Figure 1C). As for myeloid cells, classical monocyte and neutrophil concentrations were higher in treated mice, regardless of the genotype of the latter, while eosinophil counts significantly increased upon treatment only in HPV-WHIM mice. Finally, the concentrations of pDCs and of the cell population encompassing pre-DCs were increased by the X4-136 treatment in both HPV-WHIM and HPV mice (Figure 1C). Notably, in treated HPV-WHIM mice, the counts of preDCs, pDCs, classical monocytes, neutrophils, and eosinophils reached levels comparable to those observed in control HPV mice. This indicates normalization of these parameters upon CXCR4^GOF^ inhibition, despite late analysis after compound administration. Thus, the chronic oral administration of the X4-136 antagonist increases pDC and most myeloid blood cell counts in HPV-WHIM and HPV mice.

**Figure 1.**
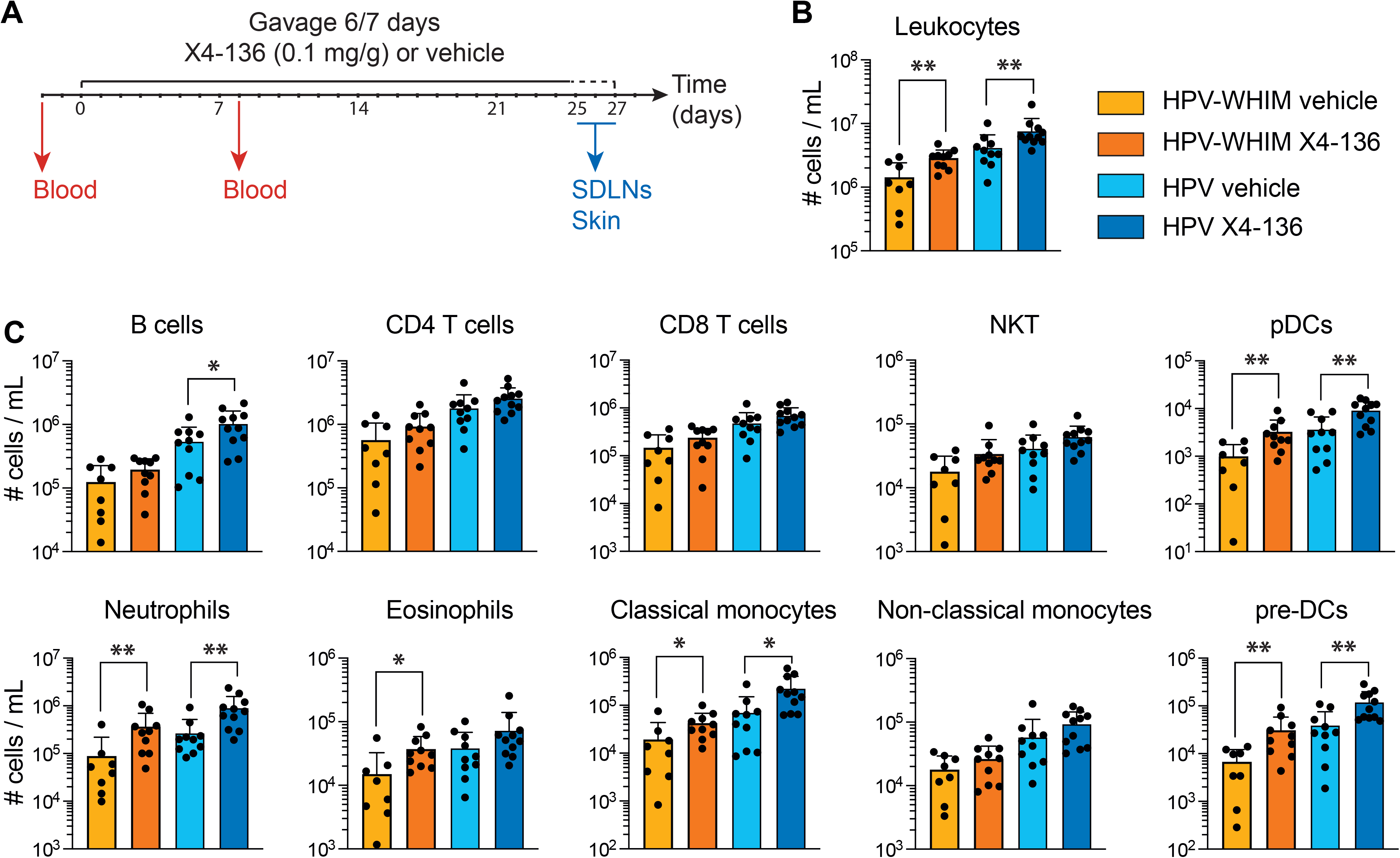
CXCR4 inhibition with the X4-136 compound normalizes leukocyte counts in HPV-WHIM mice. **(A)** Protocol design for the long-term treatment with the X4-136 compound. HPV and HPV-WHIM mice were administered 0.1 mg/g of the X4-136 compound or vehicle by gavage on 6 of 7 days for 25 to 27 days. Blood samples were collected 2 to 3 days before treatment and 8 days after treatment started, 22 hours after the last compound administration. At the end of the protocol, skin from the flanks and inguinal SDLNs were recovered. **(B)** The leukocyte and **(C)** immune population (defined as in Figure S1) concentrations were determined in the blood of HPV-WHIM and HPV mice treated or not with X4-136, after 8 days of treatment. Bar graphs show the mean ± SD. Cumulative data from 3 experiments are shown, with a total of 8-11 mice/group. Statistical analysis was performed using the two-tailed unpaired Mann-Whitney test to compare each treated group to its non-treated counterpart. \**p* < 0.05, \*\**p* < 0.01, and \*\*\**p* < 0.001.

### Inhibiting CXCR4^GOF^ with X4-136 rescues SDLN DC subset counts in HPV-WHIM mice

We next sought to determine whether and how chronic CXCR4 inhibition impacted SDLN DCs. SDLNs were recovered after 4 weeks of treatment, enzymatically dissociated, and labelled for flow cytometry analysis. MigDCs and resDCs were identified based on their expression level of CD11c and MHCII, as reported^32,40^. Type 1 and type 2 resDCs and migDCs/LCs were further identified based on the expression of the CD8α, CD11b, and CD103 markers^41^ (Figure 2A). Chronic treatment with X4-136 led to a general increase in LN cell counts in treated HPV-WHIM mice compared with vehicle-treated mice (Figure 2B). In contrast, no significant difference in LN count was observed between X4-136-treated HPV mice and their controls. As expected, the DC counts in HPV-WHIM SDLNs were lower than in their HPV counterparts, with statistical significance for pDCs, resDC1, and migDC2/LCs. Further analysis revealed that the treatment significantly increased cell counts of nearly all DC subsets in the HPV-WHIM mice, except migDC1. A similar trend was also noticed for resDCs and migDC2/LCs in treated HPV mice (Figure 2D-E). The DC activation state was then evaluated by measuring CD86 expression levels. While migDC2/LCs from HPV-WHIM mice were more activated than their controls from HPV mice in the absence of treatment, as expected^32^, this difference disappeared upon treatment (Figure S2A). Regarding CXCR4 expression, an increase in its membrane level was observed only in the treated HPV-WHIM migDC2/LCs compared to vehicle-treated ones (Figure S2B). These results support the conclusion that chronic CXCR4 inhibition by the X4-136 compound rescues DC counts in the SDLNs of HPV-WHIM mice, with phenotypic modification in migDC2/LCs.

**Figure 2.**
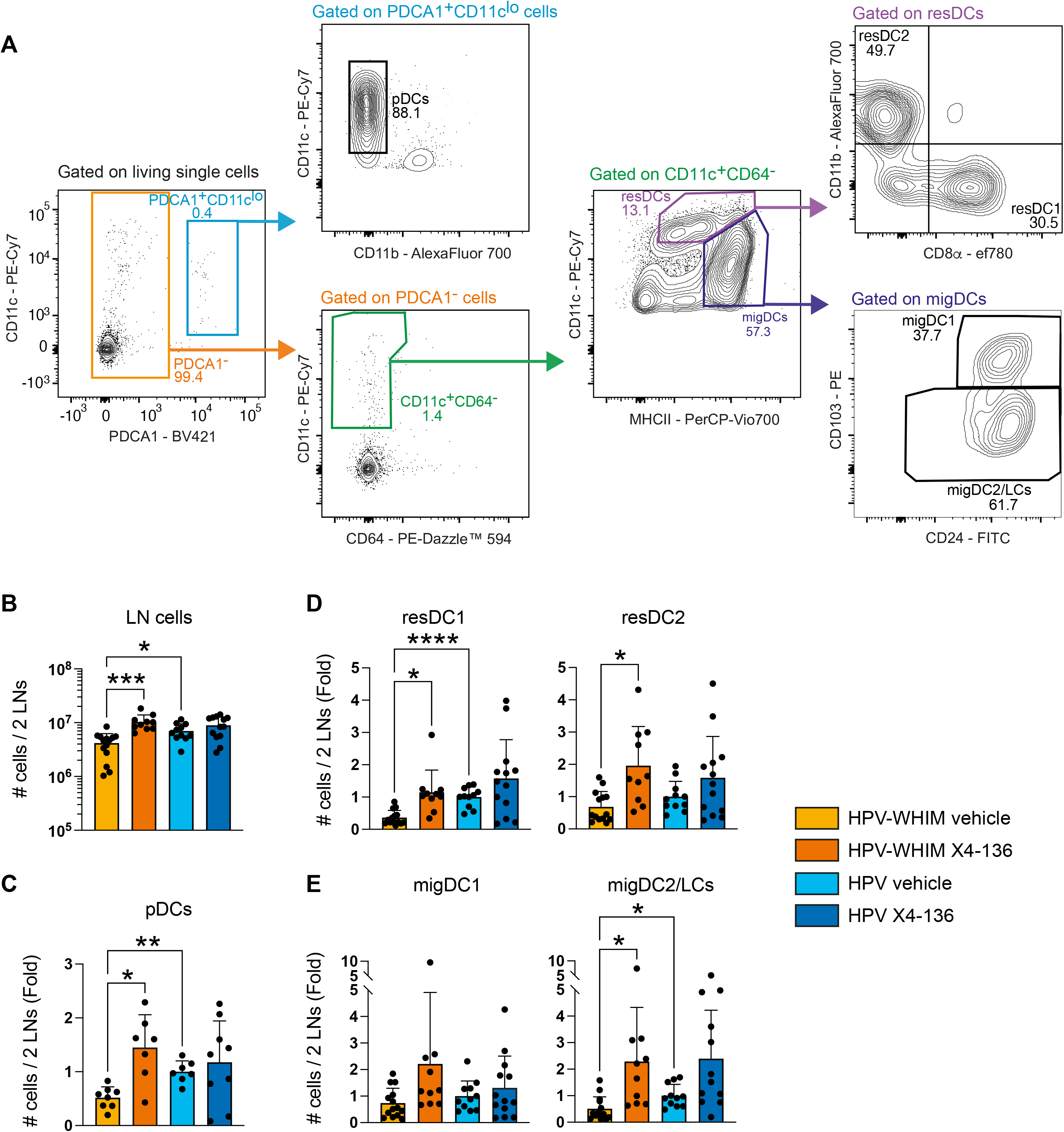
Chronic CXCR4 inhibition with the X4-136 compound rescues DC counts in the LNs of HPV-WHIM mice. **(A)** Representative contour plots show the gating strategy to identify DC subsets in inguinal lymph nodes. pDCs (PDCA1^+^CD11c^lo^CD11b^-^), resident DCs (resDCs, PDCA1^-^ CD64^-^CD11c^hi^MHCII^int to hi^), and migratory DCs (migDCs, PDCA1^-^CD64^-^CD11c^int^MHCII^hi^) were defined among viable single cells. Type 1 and 2 resDCs, resDC1 (CD8 ^+^CD11b^-^) and resDC2 (CD8 - CD11b^+^) respectively, and migDCs, migDC1 (CD103^+^) and migDC2 (CD103^-^) respectively, were further identified. **(B)** Number of cells in the inguinal lymph nodes. **(C-E)** Number of pDCs **(C)**, resDC1 and resDC2 **(D)**, and migDC1 and migDC2/LCs **(E)** in inguinal lymph nodes. Data are expressed as the fold change compared to the mean cell numbers in the HPV group for each experiment. Bar graphs show the mean ± SD for cumulative data from 2 **(C)** or 3-4 **(B, D-E)** experiments, with a total of 7-9 **(C)** or 10-14 **(B, D-E)** mice/group, respectively. Statistical analysis was performed using the Brown-Forsythe and Welch ANOVA test. \**p*< 0.05, \*\**p* < 0.01, \*\*\**p* < 0.001, and \*\*\*\**p* < 0.0001.

### Long-term treatment with X4-136 promotes the activation and relocation of type 2 dermal DCs in the skin of HPV-WHIM mice

The increase in migDC2/LC counts in the SDLNs upon chronic X4-136 treatment prompted us to assess the impact of CXCR4 inhibition on skin DC and LC populations. Therefore, mouse flank skin was recovered after 4 weeks of treatment, dissociated, and labelled for flow cytometry. Dermal DC1 and DC2 subsets, as well as LCs, were identified based on their expression levels of CD11c, CD11b, CD103, and CD24 (Figure 3A), as previously reported^32,40,41^. The counts of total skin cells were not significantly different between groups, nor were the counts of CD45^+^ cells (Figure 3B-C). The analysis of DC subset frequency among CD45^+^ cells revealed no difference either (Figure 3D). As skin DC maturation prompts their migration to SDLNs^42^, we next analyzed CD86 expression by dermal DCs and LCs. X4-136 treatment selectively promoted the activation of dermal DC2 in HPV-WHIM mice, as evidenced by a 2-fold increase in CD86 expression (Figure 3E). Such activation was not observed in the HPV group, although the compound was detected at similar levels in the plasma and skin of both treated groups (Figure S3).

**Figure 3.**
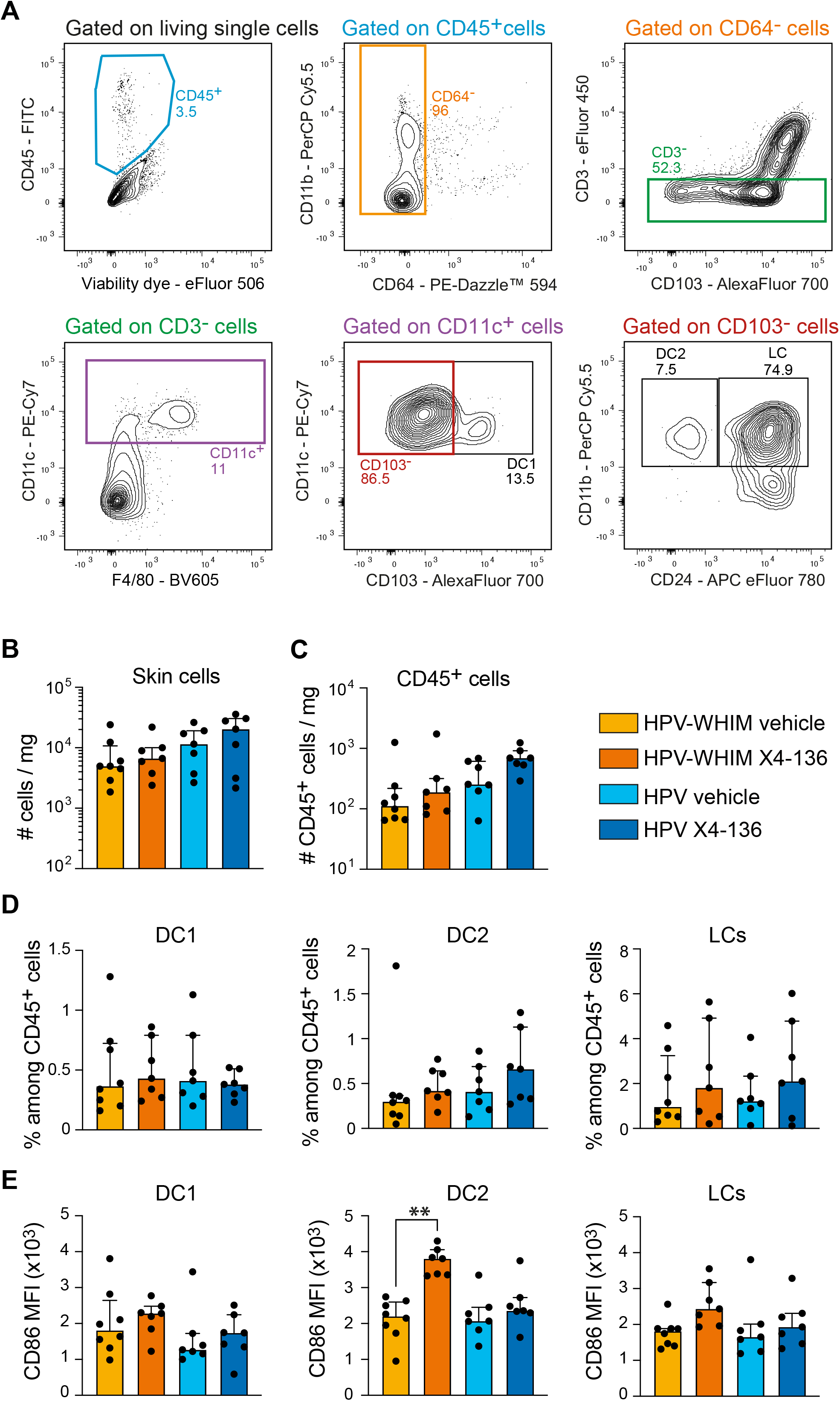
Chronic treatment with the X4-136 compound promotes dermal DC2 activation while preserving DC/LC subset counts and distribution. **(A)** Representative contour plots show the gating strategy to identify DC subsets in mouse skin. Dermal DC1 (CD11c^+^CD103^+^), DC2 (CD11c^+^CD103^-^ CD11b^+^CD24^-^), and LCs (CD11c^+^CD103^-^CD11b^+^CD24^+^) were defined among CD45^+^CD64^-^CD3^-^ viable single cells. **(B)** Number of cells per mg of skin. **(C)** Number of CD45^+^ cells per mg of skin **(D)** Percentage of DC1, DC2, and LCs among CD45^+^ skin cells. **(E)** CD86 expression by DC1, DC2, and LCs (MFI). **(B-E)** Bar graphs show the mean ± SD for cumulative data from 2 experiments, with a total of 7-8 mice/group. Statistical analysis was performed using the Kruskal-Wallis test. \*\**p* < 0.01.

To gain further insight into the effects of chronic X4-136 treatment on the local immunological landscape, we performed spatial tissue analysis on the mouse flank skin using IMC (Figure S4A) with an original panel enabling broad tissue analysis (Table S2). Unsupervised single-cell and spatial analyses were performed using the workflow described by Windhager *et al*.^38^ Dimensionality reduction and unsupervised clustering identified 15 cell clusters based on population markers expression (Table S2). Consistent with flow cytometry results, comparison of these clusters’ abundances did not reveal major differences in cell subset frequency between groups (Figure 4B). Cell phenotypes assessed by expression levels of activation or proliferation markers, such as Ki67, Granzyme B (GrzB), CD69, and MHCII (Figure S4B and S5), showed no significant impact of the treatment at the tissue level after 4 weeks. Next, to analyze the spatial distribution of cells, we performed cell-cell interaction analysis using a k-nearest neighbors’ approach and permutation testing (Figure 4C), focusing on positional changes involving fibroblasts, which are known producers of CXCL12 in the skin^43^.^43^ No significant changes in interactions between fibroblasts and myeloid cell subsets (e.g., M2 macrophages, MHCII+ macrophages, and Myeloid cells) were detected (Figure 4D). Because unsupervised data analysis failed to capture the rare DC2 subset, we conducted a supervised analysis to identify it and examine cell neighbors. Dermal DC2 from treated HPV-WHIM mice were significantly more frequently close to fibroblasts than in the untreated group (Figure 4E and 4F). Consistently, the reverse analysis of fibroblast-neighbor cells revealed an increased frequency of proximity with DC2 in treated HPV-WHIM mice (Figure 4G). This relocation clearly stood out in the global analysis landscape, confirming the quite selective effect of X4-136 on the skin DC2 compartment. Overall, our data demonstrate that treatment with X4-136 selectively promotes dermal DC2 activation and relocation toward fibroblasts in HPV-WHIM mice, likely during their migration to SDLNs.

**Figure 4.**
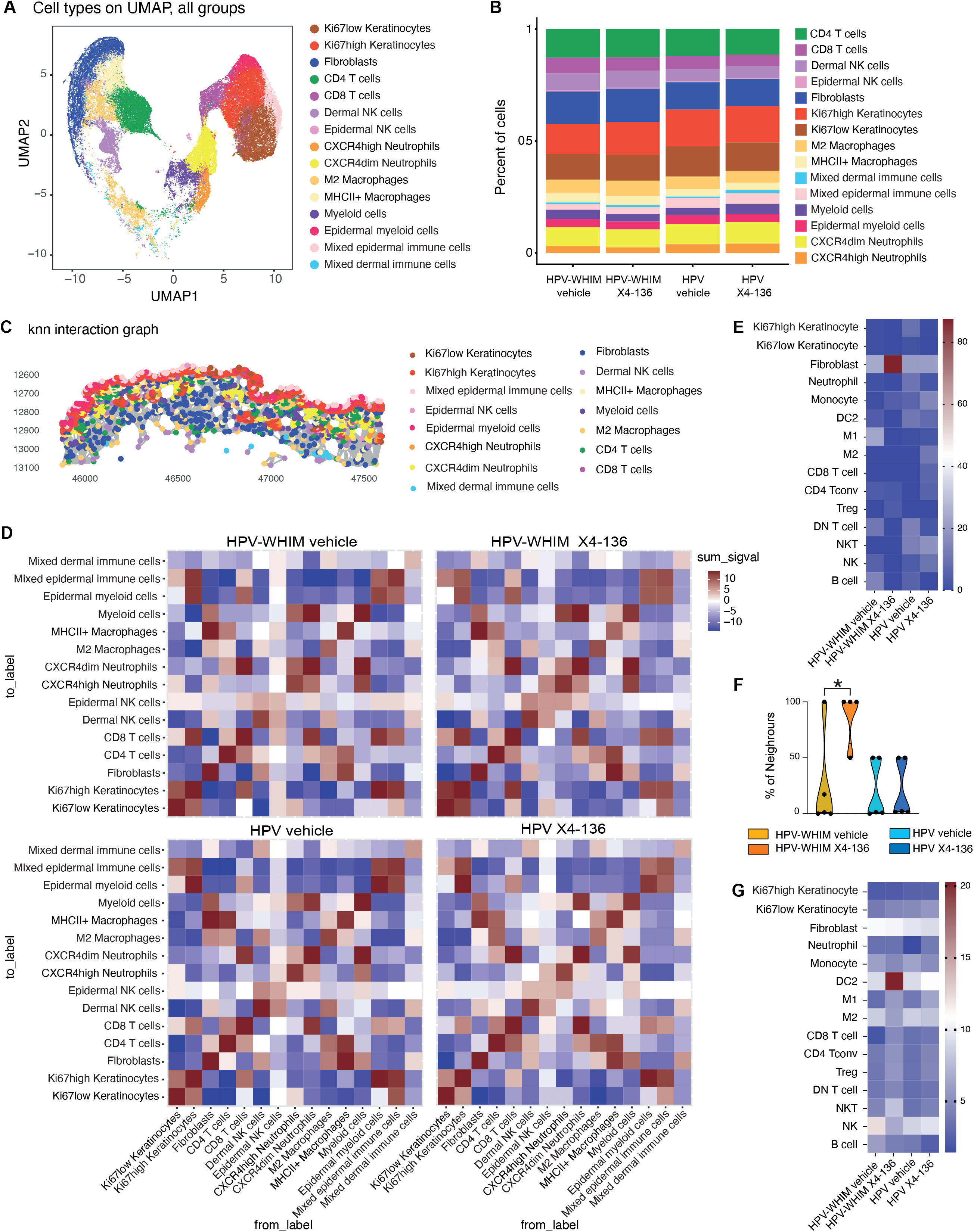
CXCR4 inhibition with the X4-136 compound promotes DC2 relocation while preserving skin architecture and immune populations. **(A)** UMAP visualization of single cells from mouse skin (pooled across all four experimental groups) colored by cell type clusters identified through unsupervised clustering. Data were acquired by imaging mass cytometry. **(B)** Bar graph shows the proportion of each cell type across experimental groups. No significant differences in cell type proportions were detected between groups. **(C, D)** Unsupervised spatial cell-cell interaction analysis using k-nearest neighbors (k=40). **(C)** Representative ROI showing spatial graph colored by cell type with k-nearest neighbor edges (grey lines) connecting spatially proximate cells. The x- and y-axis values correspond to the spatial coordinates of the pixels in the hyperspectral image (column and row indices in the image grid). **(D)** Heatmap of interaction scores from permutation testing (1000 iterations). Colors indicate: red (+1) = significant spatial attraction (p<0.01, cells interact more than expected by a null model); white (0) = no significant interaction (p>0.01); blue (-1) = significant spatial avoidance (p<0.01, cells interact less than expected by a null model). Statistical analysis did not reach significance. **(E-G)** Supervised analysis of IMC data. **(E)** DC2 were identified as CD45^+^CD3^-^NK1.1^-^ CD11c^+^MHCII^+^CD11b^+^CD103^low^ cells. Their neighboring cells were analysed by identifying cells located within 10 µm. The heatmap displays the mean percentages of interactions across different cell subsets in the different mouse groups. **(F)** Violin plots show the percentage of DC2 cells found near fibroblasts for individual mice. **(G**) Fibroblast neighboring cells. **(A-G)** Data from 2 experiments, with a total of 4-5 mice /group. Statistical analysis was performed using the Kruskal-Wallis test, with Benjamini-Hochberg p-value correction to account for multiple testing in the unsupervised analysis. \**p* < 0.05.

### CXCR4 inhibition with X4-136 promotes skin DC migration to SDLNs in WHIM mice

Increased DC counts in SDLNs from HPV-WHIM mice after chronic X4-136 treatment may result from alterations in cell proliferation, survival, and/or migration. To investigate whether X4-136-mediated inhibition promotes skin DC migration to SDLNs, we performed FITC painting experiments in WHIM mice and their WT counterparts. Mice received X4-136 or vehicle per oral gavage, and 1 hour later, a FITC solution was applied to their shaved flanks. After 22 more hours, cells from the inguinal lymph nodes were recovered and analyzed by flow cytometry to identify the FITC-positive (FITC^+^) and -negative (FITC^-^) migDC subsets coming from the painted, inflamed, skin area or from other regions, respectively (Figure 5A). The total cell counts in inguinal SDLNs from the WHIM mice remained lower than those of the WT mice despite treatment (Figure 5B). Consistent with our previous report^32^, FITC^+^ and FITC^-^ migDC counts were lower in WHIM than in WT mice in the vehicle groups (Figure 5B). In this setting, while the number of FITC^+^ migDC1 remained low upon X4-136 treatment, the FITC^+^ migDC2/LC count increased significantly in treated WHIM mice compared to untreated ones, reaching levels like those of WT mice (Figure 5C). FITC^-^ migDC1 and migDC2/LC counts also increased upon treatment in WHIM mice, although they remained lower than in untreated WT mice for migDC1 (Figure 5D). By contrast, migDC/LC counts were not increased in treated WT mice, underscoring the selective effect of the treatment on WHIM DCs. The analysis of the migDC phenotype revealed that, while WHIM migDCs from non-treated mice displayed increased CD86 levels compared to WT mice, as expected^32^, these levels were significantly reduced upon treatment in the FITC^+^ migDC2/LC and FITC^-^ migDC1 and migDC2/LC populations, albeit they remained higher than in the WT untreated controls for FITC-cells (Figure 5E-H). Consistent with the fact that these cells originate from inflamed and non-inflamed regions, respectively, CD86 levels were overall higher in FITC^+^ cells than in FITC^-^ cells. Such differences were not found in resDCs (Figure S6A). Further analysis revealed that FITC^+^ and FITC^-^ migDC1 and migDC2/LCs, as well as resDC2 from the treated WHIM mice, displayed significantly higher CXCR4 membrane levels than those from the vehicle group. Although a similar trend of increased CXCR4 expression was observed across all examined DC subsets in WT mice, statistical significance was reached exclusively in FITC^-^ migDC2/LCs (Figure 5I-L and S6B). These results demonstrate that inhibition of CXCR4^GOF^ by X4-136 enhances the migration of migDC2/LCs and migDC1 from non-inflamed tissues to SDLNs, whereas in an inflammatory context, it primarily promotes the migration of migDC2/LCs. The increase in WHIM migDC/LC migration was paralleled by a correction of their hyperactivation.

**Figure 5.**
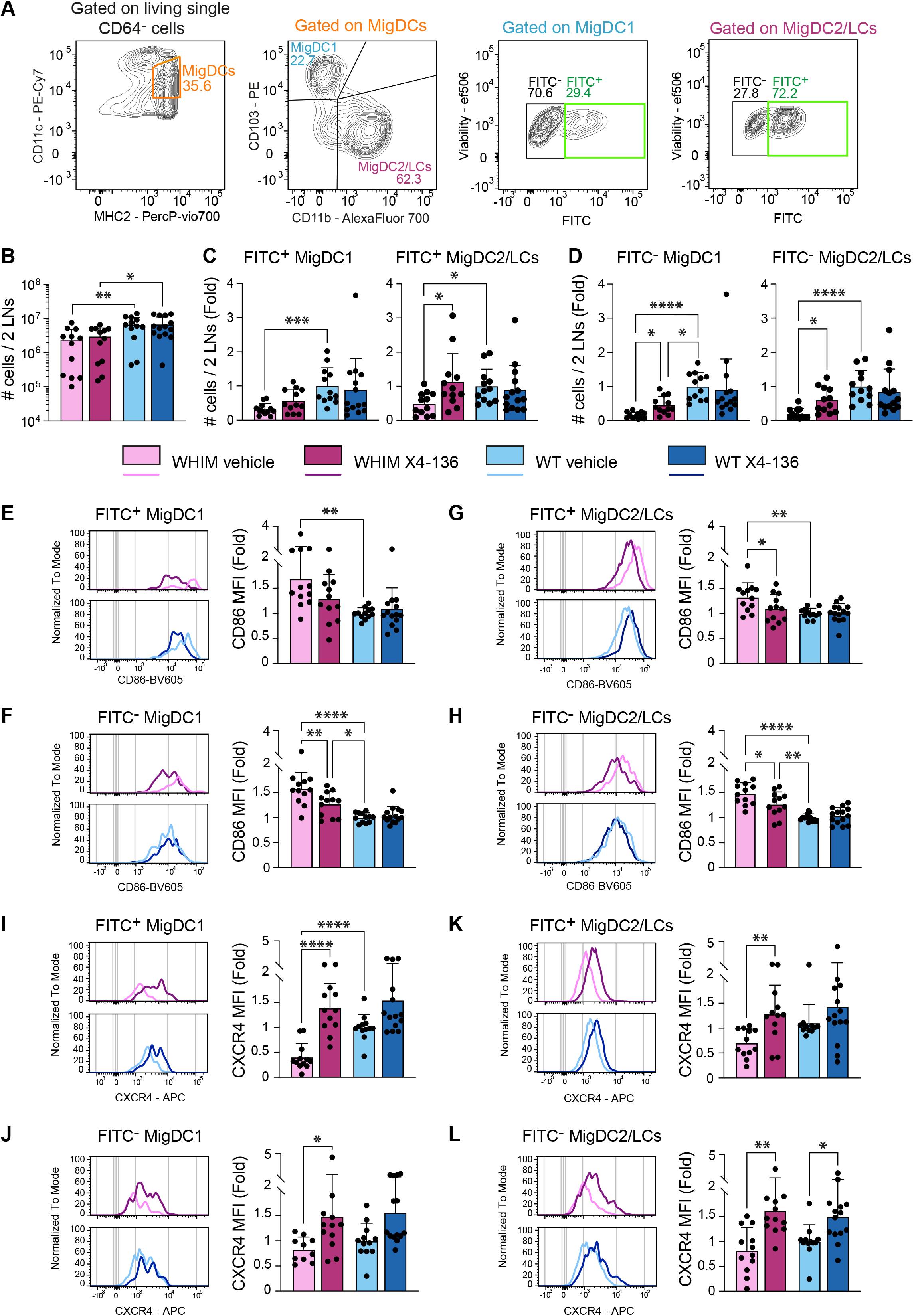
CXCR4 inhibition with the X4-136 compound promotes migDC2/LCs migration to SDLNs in the context of acute inflammation in WHIM mice. **(A)** Representative contour plots show the gating strategy to identify FITC-positive (FITC^+^) and FITC-negative (FITC^-^) migDC subsets. **(B)** Number of cells in the inguinal lymph nodes. **(C, D)** Number of FITC^+^ **(C)** and FITC^-^ **(D)** migDC1 and migDC2/LCs. **(E-H)** Bar graphs show the level of CD86 expression by FITC^+^ **(E)** and FITC^-^ **(F)** migDC1 and FITC^+^ **(G)** and FITC^-^ **(H)** migDC2/LCs. **(I-L)** Bar graphs show the level of CXCR4 expression by FITC^+^ **(I)** and FITC^-^ **(J)** migDC1 and FITC^+^ **(K)** and FITC^-^ **(L)** migDC2/LCs. **(C-L)** Data are expressed as the fold change compared to the mean cell number in the WT vehicle group for each experiment. Bar graphs show the mean ± SD for cumulative data from 5 experiments, with a total of 12-14 mice /group. Statistical analysis was performed using the Kruskal-Wallis test. \**p*< 0.05, \*\**p* < 0.01, \*\*\**p* < 0.001 and \*\*\*\**p* < 0.0001.

## Discussion

In the present study, we investigated the impact of CXCR4^GOF^ inhibition with the oral CXCR4 antagonist X4-136 on blood cell mobilization and skin DC compartment in mouse models of WHIM syndrome. We provide evidence of global rescue of WHIM DC subset defects upon X4-136 treatment, including changes in subset distribution and activation, which is anticipated to be beneficial for controlling HPV-induced lesions.

Inhibition of CXCR4 with plerixafor^30,32,45–49^ or mavorixafor^50^ promotes leukocyte mobilization across various settings, including in WHIM patients and preclinical models. We show here that CXCR4 inhibition with X4-136 also normalizes circulating myeloid leukocyte and pDC counts in HPV-WHIM mice. This suggests that X4-136-based CXCR4 inhibition efficiently unwinds CXCL12-CXCR4 interactions in the BM, and possibly in other tissues, thereby promoting leukocyte mobilization. Specifically, we observed an increase in blood pre-DC and pDC concentrations in treated HPV-WHIM mice, reaching levels in control HPV mice, suggesting that the normalization of the SDLN pDC, resDC1, and resDC2 counts would result from the replenishment of these populations. However, despite elevated circulating pre-DC and pDC concentrations, there was no increase in SDLN pDC, resDC1, or resDC2 counts in HPV mice. This suggests that additional checkpoints, likely linked to Flt3L and niche availability^51^, control pDC entry and pre-DC entry and differentiation in SDLNs.

We explored the impact of long-term CXCR4 inhibition on skin DCs and migDC subsets. Deep unsupervised profiling of the cutaneous immune landscape by IMC demonstrated only discrete differences in skin immune cells between X4-136-treated and untreated HPV and HPV-WHIM mice, indicating that tissue architecture and cellular composition are preserved despite chronic CXCR4 inhibition. However, assessing the position and activation of dermal DC2, the subset that displays normalized migration to SDLNs in treated HPV-WHIM, revealed that they relocated near fibroblasts and showed a 2-fold increase in their expression of the CD86 activation marker in treated HPV-WHIM compared with their controls, a difference not observed in the HPV-treated control group. CXCL12 is mainly produced by reticular fibroblast subsets adjacent to lymphatics, making the redistribution of DC2s toward fibroblasts after CXCR4^GOF^ inhibition seem paradoxical. Still, recent single-cell analyses in human skin have identified fibroblastic reticular cell-like fibroblasts in perivascular regions that, in addition to CXCL12, express cues supporting DC migration, such as CCL19 and CH25H-derived oxysterols^52^, and engage in direct interactions with migrating DCs^53^. While this fibroblast subset is rare in healthy mouse skin, oxysterol-producing fibroblasts have been reported during chronic skin inflammation in lupus mouse models^54^. Alternatively, the proximity of DC2 cells to fibroblasts in treated HPV-WHIM mice could result from their recruitment to nearby lymphatic vessels, as these cells would become able to respond to CCL19/CCL21 chemotactic signals produced by lymphatic endothelial cells upon CXCR4^GOF^ inhibition. The selective DC2 relocation within the skin may favor upregulation of CD86 expression in these cells, which would explain the selective effect of X4-136 on this DC subset, thereby contributing to their improved egress from the skin and migration to SDLNs and the normalization of migDC2/LC counts in HPV-WHIM SDLNs to levels like those in treated HPV mice.

Whether CXCR4^GOF^ inhibition modified skin DC migration to SDLNs was directly assessed by tracking skin DC migration in FITC-painting experiments, after a single X4-136 administration. In this setting, where cell proliferation and survival have minimal impact on DC counts, inhibiting CXCR4^GOF^ primarily increased the number of FITC^+^ migDC2/LCs, indicating enhanced migration. Based on the known migration kinetics of LCs and dermal DCs^55^, this increase is most probably accounted for by DC2 rather than LCs. These results support the conclusion that, in inflammatory contexts, whether acute upon FITC-painting or chronic in HPV transgenic mice, CXCR4^GOF^ inhibition promotes dermal DC2, but not DC1, migration to SDLNs. FITC-painting experiments further revealed increased FITC^-^ migDC1 and migDC2/LC counts in the SDLNs of WHIM mice. These data support the conclusion that the inhibition with X4-136 promotes the migration of both dermal DC1 and DC2 from non-inflamed tissues. This suggests a scenario in which DC2 would be more responsive than DC1 to local CXCL12 levels in inflamed dermis, possibly due to lower CCR7 expression^32^. Taken together, our results support the conclusion that CXCR4 inhibition with X4-136 restores dermal DC migration to SDLNs in the WHIM context, as well as pDC and resDC entry and/or differentiation in these lymphoid organs. Given the known importance of coordinated action among DC subsets in fighting viral infection^56–59^, such normalization of the whole DC compartment in SDLNs could be instrumental in unleashing efficient antiviral T-cell responses in the WHIM context. Testing this hypothesis warrants additional dedicated studies.

Boosted dermal DC migration to SDLNs upon CXCR4^GOF^ inhibition was associated with reduced CD86 expression levels in migDCs collected from SDLNs, witnessing a normalization of their activation. While this may appear contradictory to the finding that CXCR4^GOF^ inhibition promoted dermal DC2 activation in chronic inflammation, these results must be considered in light of the much higher activation levels of migDCs in SDLNs than in skin DCs (Figure 6). The normalization of CD86 expression was observed selectively for the migDC subsets whose migration was increased, regardless of whether they originated from inflamed or non-inflamed tissues. This supports the conclusion that disrupting CXCR4-CXCL12 interactions in the dermis of WHIM mice results in the arrival of less activated migDCs into the SDLNs by facilitating the egress of skin DCs from the CXCL12-rich dermis. In contrast, in untreated WHIM mice, only highly activated DCs would succeed in exiting the dermis. Such an effect would not occur in the WT context, where the normal desensitization process naturally frees dermal DCs from the CXCL12-driven attraction. The increase in CXCR4 membrane levels observed in migDCs upon X4-136 administration may result from reduced receptor internalization or resensitization, as a direct consequence of X4-136 binding to CXCR4 or due to reduced CXCL12-mediated desensitization. It may alternatively reflect the release of DCs with high CXCR4 membrane expression, which are otherwise trapped in the dermis or lymphatic vessels of WHIM mice due to CXCL12-dependent retention. The CXCR4 upregulation in FITC^-^ migDC2 from treated WT mice corroborates a model in which CXCL12-rich niches regulate receptor availability and DC migratory capacity.

**Figure 6.**
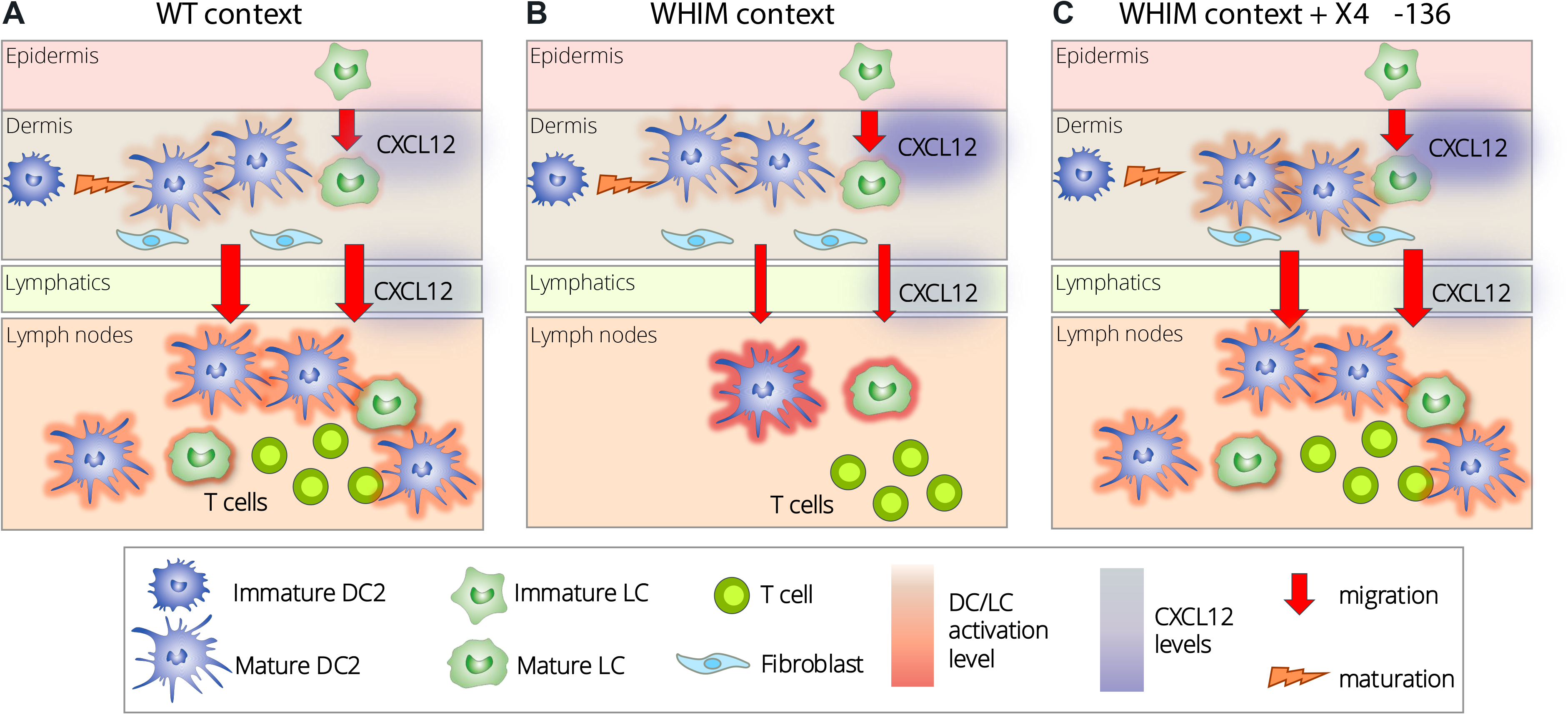
Inhibition of CXCR4^GOF^ by X4-136 reshapes skin DC2 location, activation and migration to SDLNs. **(A)** In WT contexts, dermal DC2 migrate homeostatically toward SDLNs (thick red arrow) as they become activated (light red halo around cells). Some of them are detected close to fibroblasts. **(B)** In WHIM contexts, dermal DC2 show reduced migration to the SDLNs (thin red arrow), and the few migDC2 recovered in these lymphoid organs are hyperactivated (dark red halo around cells). DC2 are not found close to fibroblasts. This could be linked to previously reported higher CXCL12 levels in the skin^60^. **(C)** Upon CXCR4^GOF^ inhibition with X4-136 in WHIM contexts, dermal DC migration to SDLNs is restored (thick red arrow), and migDC2 are not overactivated (medium reddish hallow around cells). This phenomenon is associated with the relocation of dermal WHIM DC2 close to fibroblasts in the skin and their intermediate activation (medium-red halo around cells). The potential impact on CXCL12 levels is not known.

The more pronounced effect of X4-136 on the skin DC compartment in the WHIM context is unlikely to result from differential binding or antagonism of wild-type and mutant CXCR4, as this compound inhibits calcium mobilization similarly in cells expressing each receptor form. We also excluded the hypothesis that X4-136 was distributed differently in the blood and skin of HPV-WHIM and HPV mice. In addition, circulating pre-DC and pDC levels were similarly increased in the HPV and HPV-WHIM mice, suggesting that environmental modifications may explain the selective effects in the WHIM context. Previously, we reported that CXCL12 levels were increased in the skin of WHIM mice compared with their controls^60^, which may contribute to the differences observed here. In addition, an increase in local inflammation in HPV-WHIM mice compared to HPV mice^31,32^, combined with differences in DC niche availability in the context of CXCR4^GOF^, as reported for B cells^61^, is likely a key factor contributing to the alterations in the DC compartment.

Finally, short-term CXCR4 inhibition with X4-136 did not impair DC migration, unlike acute administration of motixafortide in a WT context or AMD3100 in both WHIM and WT contexts. Differences in both the compounds’ pharmacological properties and their administration route likely contribute to these distinct treatment outcomes. While X4-136 is observed to have a longer half-life than AMD3100, the latter exhibits a slower off-rate, suggesting it may have an insurmountable mode of action. Combined with the administration mode, this may account for differences in bioavailability, pharmacology, body distribution, and ultimately, biological effects between the two antagonists. Along this line, a previous study comparing AMD3100 and KRH3955, which bind to distinct regions of CXCR4, reported distinct mechanisms of hematopoietic progenitor cell (HPC) mobilization.^62^ AMD3100’s action depended on its capacity to reverse the CXCL12 chemokine gradient at the BM endothelium, whereas KRH3955 promoted HPC egress from the BM via a more direct mechanism.^62,63^ At the level of dermal lymphatic vessels, AMD3100 may not be able to shape a mobilizing CXCL12 gradient, thereby failing to promote skin DC migration to SDLNs, as opposed to CXCR4 antagonists acting differently. Further mechanistic exploration of the effects of the CXCR4 antagonists on the DC compartment is needed to clarify these issues.

In conclusion, our work suggests that the functional restoration of the DC compartment can be achieved in the WHIM context through chronic CXCR4^GOF^ inhibition and supports the hypothesis that this may rewire skin immunity. Thus, the beneficial effects of plerixafor and mavorixafor on HPV mucocutaneous lesions in WHIM syndrome patients^14,18–20,22^ could result from rescuing DC functions. Future work is required to explore whether the improvement of DC migration to SDLNs we report would translate into more efficient anti-HPV T-cell responses.

## Acknowledgments

The authors thank Marie-Laure Aknin from the CYM core facility of the “Unité Mixte de Service, Ingénierie et Plateformes au Service de l’Innovation Thérapeutique (UMS-IPSIT) for access to its equipment and infrastructures and acknowledge Cristian Dicu from the ANIMEX core facility from the UMS-IPSIT and Laura Derout for excellent technical help. They also thank the Région Ile-de-France and INSERM for financial support to the CYM and ANIMEX core facilities. They acknowledge the cytometry flow core facility Hyperion (Brest, France) for its technical assistance, as well as the European FEDER grant program Progos RU 000950 members of the Scientific Interest Group Biogenouest.

X4 Pharmaceuticals, Inc. provided financial support for the salary of A.O., as well as the X4-136 compound, and support for mouse breeding and housing and antibody purchase. This work was also supported by grants from Marie Skłodowska-Curie actions Innovative Training Networks (ITN) “ONCOgenic Receptor Network of Excellence and Training 2.0” (FB and GSL: http://www.oncornet.eu/) and Fondation pour la Recherche Médicale (EQU202203014751 Team Project; https://www.frm.org/). This work was supported as part of the France 2030 programme “ANR-11-IDEX-0003”, from the GS HeaDS of the Université Paris-Saclay. It was also supported by the MetaboHUB infrastructure funded by the Agence Nationale de la Recherche under the France 2030 program (MetaboHUB ANR-11-INBS-0010; MetEx+ ANR-21-ESRE-0035; MetaboHUB (JVCE) ANR-24-INBS-0012). M.D. was supported by a doctoral fellowship from the Doctoral School “Innovation Thérapeutique du Fondamental à l’Appliqué” (ED 569). G.C.M. was supported by the Marie Skłodowska-Curie actions Innovative Training Networks (ITN) “ONCOgenic Receptor Network of Excellence and Training 2.0”.

## Authorship Contributions

A.O., A.J.R., M.D, S.S.V, M.O., F.M.N., Y.D., A.L.N, A.M., A.P., K.K., K.Z., F.B. and G.S.L. designed and performed experiments. A.O., M.P., S.S.V., A.L.N., A.M., A.P., P.H., F.B. and GSL analyzed the results. A.O., M.P., A.J.R., M.D, G.C.M, F.B. and G.S.L. wrote the original draft. A.O., M.P., A.J.R., M.D, G.C.M., K.Z., L.D, S.Z., C.H.N., R.J., A.T., C.D., F.B. and G.S.L. critically discussed the results, validated the conclusions and reviewed the manuscript. F.B. and G.S.L. acquired funding and supervised the study.

