## Supplemental material for "CXCR4 antagonism restores dendritic cell migration and activation in a WHIM syndrome mouse model"

#### Supplemental Methods

##### Ligand binding inhibition assay and washout experiments to determine residence time

Jurkat cells were washed with assay buffer (HBSS + 20 mM HEPES buffer + 0.2% bovine serum albumin, pH 7.4) and incubated for 15 min at room temperature with different concentrations of X4-136 compound diluted in assay buffer. Subsequently, human CXCL12-AlexaFluor647 (26 ng/mL, Almac, Craigavon, United Kingdom) was added to the compound-preincubated cells. The cells were incubated for 30 min at room temperature. Thereafter, the cells were washed twice in assay buffer, fixed with 1% paraformaldehyde in PBS, and analyzed by flow cytometry (CytoFlex, Beckman Coulter). For washout experiments, Jurkat cells were pre-incubated with the X4-136 compound at IC90 concentration (80 nM), washed, and human CXCL12-AlexaFluor647 was added at various time points. After a 30-min incubation, cells were washed and processed as previously described.(1) Mean fluorescence intensity (MFI) of CXCL12-AF647 was determined (FCS express software). The percentage of inhibition was calculated according to the formula: $[1-((MFI-MFI_{NC})/(MFI_{PC}-MFI_{NC}))]*100$ where MFI is the MFI of cells in the presence of an inhibitor, $MFI_{NC}$ is MFI of cells in the absence of the ligand and $MFI_{PC}$ is MFI of cells in the presence of the ligand alone.

##### Calcium mobilization assay

Stable K562 clones expressing the WT or S338X CXCR4 ($10^5$ cells/well) were seeded in black 96-well plates with transparent bottom coated with poly-L-lysine (BioCoat®, Corning) and serum-starved for 24 h. The medium was removed, and cells were loaded with 100 µL of fluo-4 AM (3 µM, Invitrogen) dye solution for 45 min at 37 °C. The dye solution was prepared by diluting fluo-4 powder in DMSO + 10% Pluronic F127 (Thermo Fisher Scientific, Waltham, Massachusetts, USA) to 3 mM and then preparing the final dilution in assay buffer (HBSS with $Ca^{2+}$ and $Mg^{2+}$, 20 mM HEPES, 0.375 g/L NaHCO3, 0.1% BSA + 0.77 g/L probenecid [Thermo Fisher Scientific]). Subsequently, 100 µL of X4-136 dilutions or assay buffer alone were added, and the plates were equilibrated in the plate reader for an additional 20 min at 37 °C. CXCL12 was injected with simultaneous measurement of fluorescent signal (FlexStation® 3 Multi-Mode Microplate Reader; Molecular Devices, San Jose, California, USA). Raw traces were analyzed in SoftMax Pro 7 Software (Molecular Devices). The arbitrary units were calculated as the difference between maximal and minimal signal after treatment injection, normalized to the baseline signal before injection. Maximal effect ($E_{max}$) and half-maximal effective and inhibitory concentrations ($EC_{50}$ and $IC_{50}$, respectively) were calculated in Prism (GraphPad Software, San Diego, California, USA).

**Absolute quantification of X4-136 by liquid chromatography coupled with mass spectrometry (LC-MS/MS)**

All solvents (VWR France) were LC/MS grade.

*X4-136 extraction*

Ear biopsy samples were weighed precisely (20-100 mg) into Precellys tubes (MK28-R, Bertin Technologies, Montigny-le-Bretonneux, France). Five volumes of a solution of Bovine serum albumin (BSA, 30 g/L) in water/ACN (50/50, v/v) were added and ear biopsies were lysed using a Precellys device (Precellys Evolution, Bertin Technologies, Montigny-le-Bretonneux, France) for 3 x 30 s – 6500 rpm – 15 s – 5°C. Protein precipitation was performed by adding 300 µL of acetonitrile, 1% of NH3OH. Samples were then vortexed for 15 s and centrifuged at 20,000 g for 10 min at 5°C to remove cell debris and precipitate proteins. Supernatants were then dried under a stream of nitrogen at 40°C using a TurboVap instrument (Biotage, UK). Prior to LC-MSMS analysis, dried extracts were resuspended with 100 µL of ammonium formate 25 mM, and 0.2% HCOOH/Acetonitrile, and 0.1% HCOOH (95/5, v/v). After a final centrifugation step at 20,000 g for 10 min at 4°C, supernatants were recovered and transferred into 0.3 mL vials.

*Preparation of calibration solutions (CS) and quality controls (QCs)*

Calibration solutions were performed in plasma or in a solution of BSA (30 g/L in water/ACN (50/50, v/v) to quantify plasma and ear homogenate respectively. The calibration curve ranged from 1.00 to 400 ng/mL for plasma and 0.39 ng/mL to 200 ng/mL for BSA solution. QCs were performed in plasma or in ear biopsy lysate. QC concentrations ranged from 4.00 to 340 ng/mL for plasma and 2.50 ng/mL to 160 ng/mL for ear homogenate. CS and QCs were extracted and treated as samples.

*LC-MS/MS analysis*

Targeted LC-MS/MS measurements were performed using an ACQUITY UPLC® System coupled to a XEVO™ TQ-XS Mass Spectrometer from Waters (Guyancourt, France). The software interface was Masslynx (version 4.2) (Waters, Guyancourt, France). The LC separation was performed on a BEH C18 2.1*50 mm column (Waters, Guyancourt, France) maintained at 40°C. Mobile phase A consisted of an aqueous buffer of 25 mM ammonium formate, 0.2% formic acid, and mobile phase B of 100% acetonitrile, 0.1% formic acid. Chromatographic elution was achieved with a flow rate of 0.6 mL/min. After injection of 5 µL of the sample, the gradient conditions were maintained up to 1.0 min at 5% B, ramped from 5% B to 50% B between 1.0 and 2.0 min, from 50% B to 100% B between 2.0 and 2.01 min and was maintained up to 2.50 min at 100% B. The gradient was then returned to its initial conditions between 2.5 and 2.51 minutes and was maintained during an equilibration period prior to the next injection. The column effluent was directly introduced into the electrospray source of the mass spectrometer, and analyses were performed in positive ion electrospray multiple reaction monitoring mode. Multiple transitions monitored were m/z 339.2 > 267.9 for X4-136. In these conditions, the mean X4-136 retention time was 1.74 min.

*Data processing and quantification*

Masslynx software was used for peak detection and integration. For ear biopsy, concentrations were normalized with the exact mass weighed. Quantification was performed using linear regression with

$1/X^2$ weighing and calibration ranges from 1.00 to 400 ng/mL for plasma and 2.35 to 1200 ng/g for ear biopsy. Quantification was carried out using a standard calibration curve.

**Mouse sample processing**

Blood was collected in a tube containing 10 µL of EDTA 0.5 M (Thermo Fischer Scientific, France). Plasma was recovered, and red blood cells were lysed with ammonium-chloride-potassium buffer. For skin cell isolation, tissue pieces from the flanks were incubated for 90 min in dispase (2.5 U/mL, Gibco) at 37°C. The epidermis and dermis were dissociated and cut into small pieces that were further incubated for 1 h in collagenase D (1 mg/mL, Roche) and DNase I (12.5 U/mL, Roche) at 37°C. Tissue dissociation was completed using 18-G needle syringes, and cell suspensions were further incubated for 15 min at 37°C and filtered with 70 µm cell strainers. For SDLN cell isolation, inguinal LNs were harvested, perfused, and incubated with collagenase D (1 mg/mL, Roche) and DNase I (12.5 U/mL, Roche) at 37°C for 30 min. LN suspensions were filtered with 70 µm cell strainers.

**Flow cytometry**

Cell suspensions ($10^7$ to $5.10^7$ cells/mL) in PBS-fetal bovine serum (FBS) 2% were used. Fc receptors were blocked with anti-CD16/32 monoclonal antibodies (mAbs) for 5 min at 4°C. The viability dye and labeled mAbs were then added (Supplementary Table 1), and cells were incubated for 25 min at 4°C. After 3 washes in PBS-FBS 2%, cells were fixed with BD Cytofix (BD Biosciences, USA) for 20 min at 4°C and washed once before acquisition. Data were acquired using an LSR Fortessa (BD Biosciences, USA) and analyzed with FlowJo Software (Tree Star Inc., USA). Fluorescence-minus-one (FMO) and isotype control (for CXCR4) controls were used to determine the positivity thresholds.

**Imaging Mass Cytometry**

*Staining*

Carrier-free antibodies were conjugated to metal tags using the MaxPar® labeling kit (Fluidigm) following the manufacturer's instructions. Formalin-fixed, paraffin-embedded (FFPE) skin biopsies were cut into 4 µm slices, deparaffinized, rehydrated, and incubated in antigen retrieval solution (EDTA buffer, pH 9) at 96°C for 20 minutes. The slides were cooled to room temperature (RT), rinsed in double-distilled water ($ddH_2O$) and then in Tris-buffered saline (TBS). To block nonspecific binding, samples were pre-incubated in 3% bovine serum albumin (BSA) in PBS for 30 minutes at RT. After removing the blocking solution, 100 µL of the antibody mix was applied to each section. Slides were incubated overnight at 4°C in a humidified chamber. The following day, sections were washed twice in TBS, stained with Intercalator-Ir diluted in TBS for 5 minutes at RT in a humidified chamber, and rinsed in TBS for 5 minutes and in $ddH_2O$ for 10 seconds. Finally, the slides were air-dried at 37°C for 20 minutes before acquisition on a Hyperion imaging system coupled to a Helios Mass Cytometer (Standard BioTools), at a laser frequency of 200 Hz and laser power of 3 dB.

*Analysis*

Data were acquired on a Hyperion imaging system coupled to a Helios Mass Cytometer (Standard BioTools), at a laser frequency of 200 Hz and laser power of 3 dB. For each recorded region of interest (ROI), stacks of 16-bit single-channel TIFF files were exported from MCD binary files using MCD™ Viewer 1.0 (Standard BioTools). Cell-based morphological segmentation was carried out by using

Visiopharm software (Visiopharm). To extract quantitative data, CVS files were uploaded and analysed with OMIQ software. The neighborhood microenvironment to characterize cell-cell interactions in tissue samples was performed using custom Python (Pretreatment imaging mass cytometry reveals spatial immune architecture associated with checkpoint inhibitor benefit in metastatic cutaneous melanoma, under review in JITC). For one mouse in the treated HPV-WHIM group, only one DC was detected by supervised gating; this mouse was removed from the neighboring analysis. Unsupervised analysis was performed with the imcRtools using R versions 4.5.1 and 4.5.2 in RStudio. Expression data were arcsinh-transformed (cofactor = 1) and analyzed following the Windhager *et al*. workflow for multiplexed image analysis(2). Dimensionality reduction was performed using Uniform Manifold Approximation and Projection (UMAP) in the scatter package based on the following population markers: ColT1, PanKerat, CD45, ECad, B220, F4_80, CD103, CCR2, TCRgd, CD11c, NK1.1, Ly6G, CD4, CD3, CD207, and CD8. Batch correction was applied using Harmony to account for inter-sample variability. Unsupervised clustering was conducted using the Rphenograph algorithm (k=75). Spatial graphs were constructed using k-nearest neighbors (k=40), and cell-cell interaction testing was performed with the testInteractions function from imcRtools. Data visualization was performed using the dittoSeq R package.

### Supplemental Figures

A

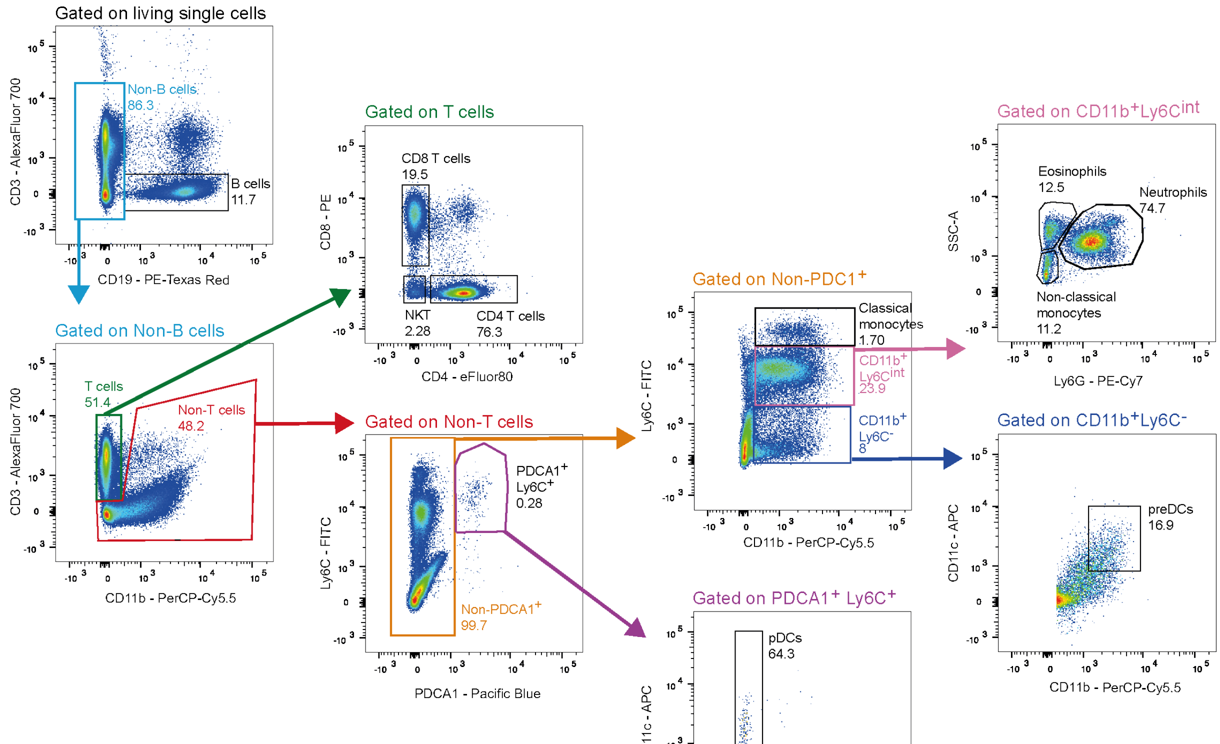


B

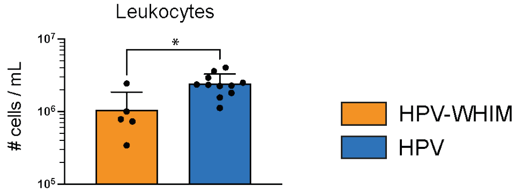


C

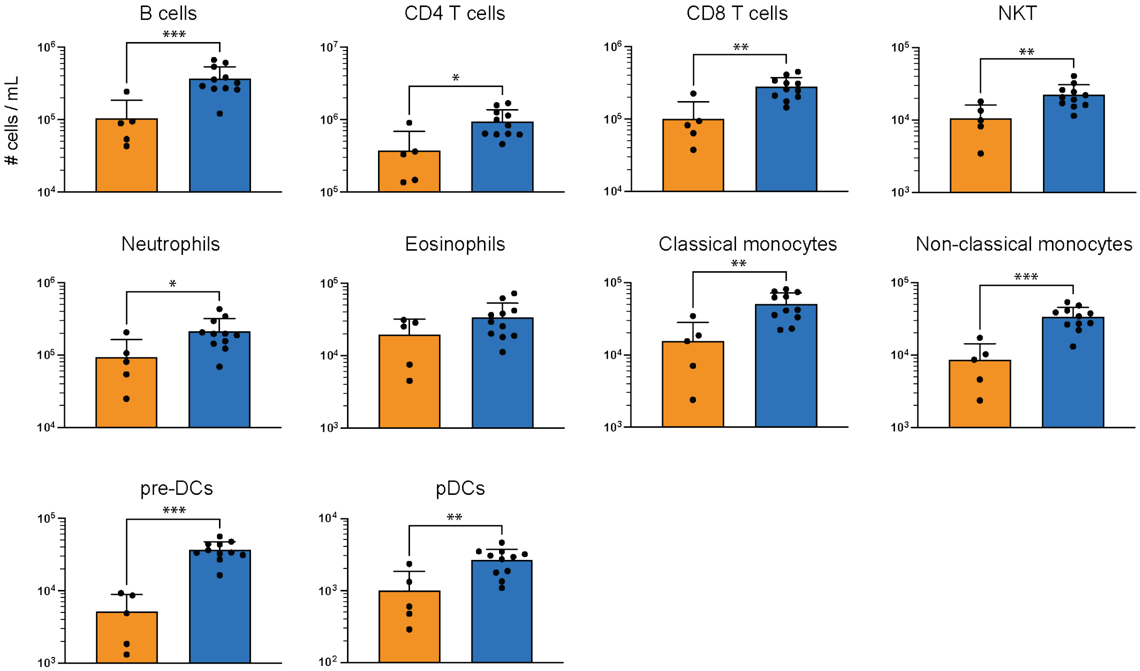

**Supplemental Figure 1: HPV-WHIM mice display marked leukopenia. (A)** Representative dot plots show the gating strategy to define blood B cells ($CD19^{+}$), CD4 ($CD3^{+}CD4^{+}CD8^{-}$) and CD8 ($CD3^{+}CD4^{-}CD8^{+}$) T cells, NKT cells ($CD3^{+}CD4^{-}CD8^{-}$), pDCs ($CD19^{-}CD3^{-}PDCA1^{+}Ly6C^{+}CD11c^{lo}CD11b^{-}$), classical monocytes ($CD19^{-}CD3^{-}PDCA1^{-}Ly6C^{hi}CD11b^{+}$), non-classical monocytes ($CD19^{-}CD3^{-}PDCA1^{-}Ly6C^{int}CD11b^{+}SSC\text{-}A^{lo}Ly6G^{-}$), neutrophils ($CD19^{-}CD3^{-}PDCA1^{-}Ly6C^{int}CD11b^{+}SSC\text{-}A^{hi}Ly6G^{+}$), eosinophils ($CD19^{-}CD3^{-}PDCA1^{-}Ly6C^{int}CD11b^{+}SSC\text{-}A^{hi}Ly6G^{-}$), and the gate containing pre-DCs ($CD19^{-}CD3^{-}PDCA1^{-}Ly6C^{-}CD11b^{+}CD11c^{+}$) among viable single cells. **(B)** The total leukocyte and **(C)** immune cell population concentrations were determined in HPV-WHIM and HPV mice. Bar graphs show mean ± SD. N=5-11 mice/group. Data are from one representative experiment out of 3. Statistical analysis was performed using the two-tailed unpaired Mann-Whitney test. *$p < 0.05$, **$p < 0.01$ and ***$p < 0.001$.

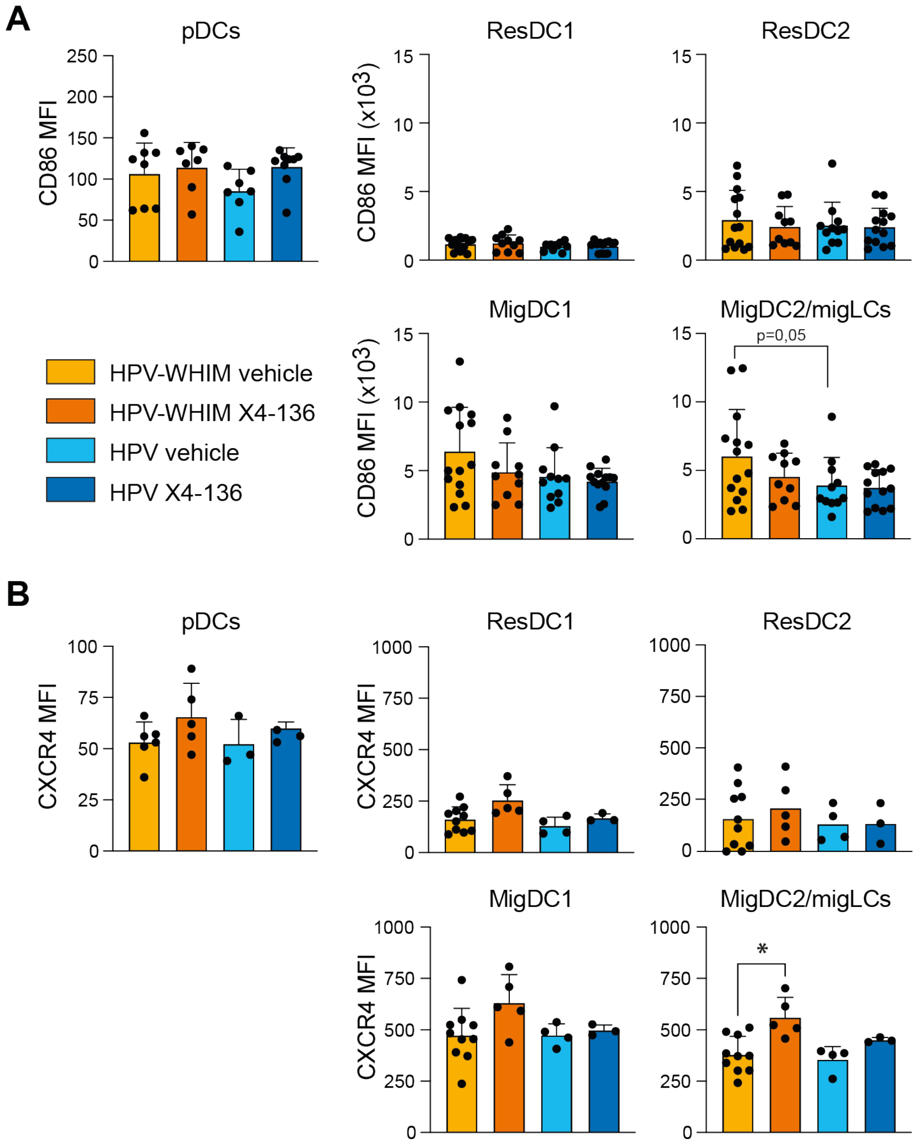


**Supplemental Figure 2. Chronic CXCR4 inhibition with the X4-136 compound does not modify the activation profile of SDLN DCs but increases CXCR4 membrane expression in MigDC2/LCs.** CD86 **(A)** and CXCR4 **(B)** expression were assessed in pDCs, resDC1, resDC2, migDC1, and migDC2/migLCs. Bar graphs show the mean MFI ± SD for cumulative data from 2-4 **(A)** or 1-2 **(B)** experiments, with 7-14 **(A)** or 3-10 **(B)** mice/group. Statistical analysis was performed using one-way Anova **(A)** and the Kruskal-Wallis test **(B)**.

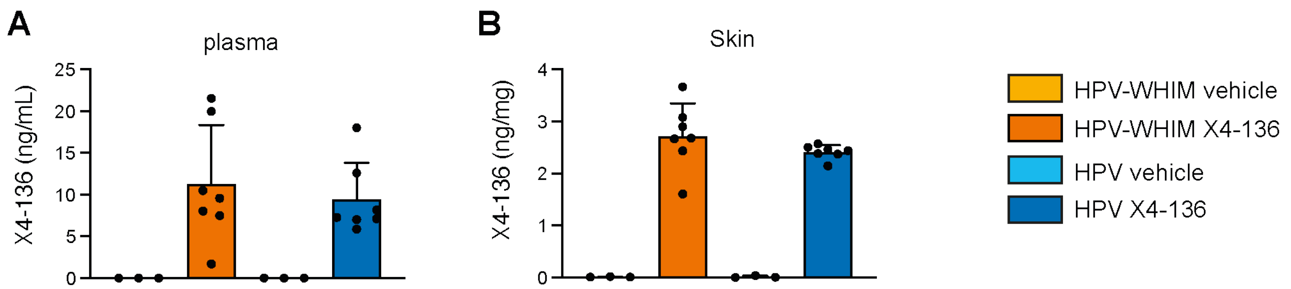


**Supplemental Figure 3. Absolute quantification of the X4-136 compound in the plasma and skin of treated mice.** The X4-136 compound was detected in the **(A)** plasma and **(B)** skin of mice after 25-27 days of treatment. Bar graphs show the mean ± SD for data from 2 experiments.

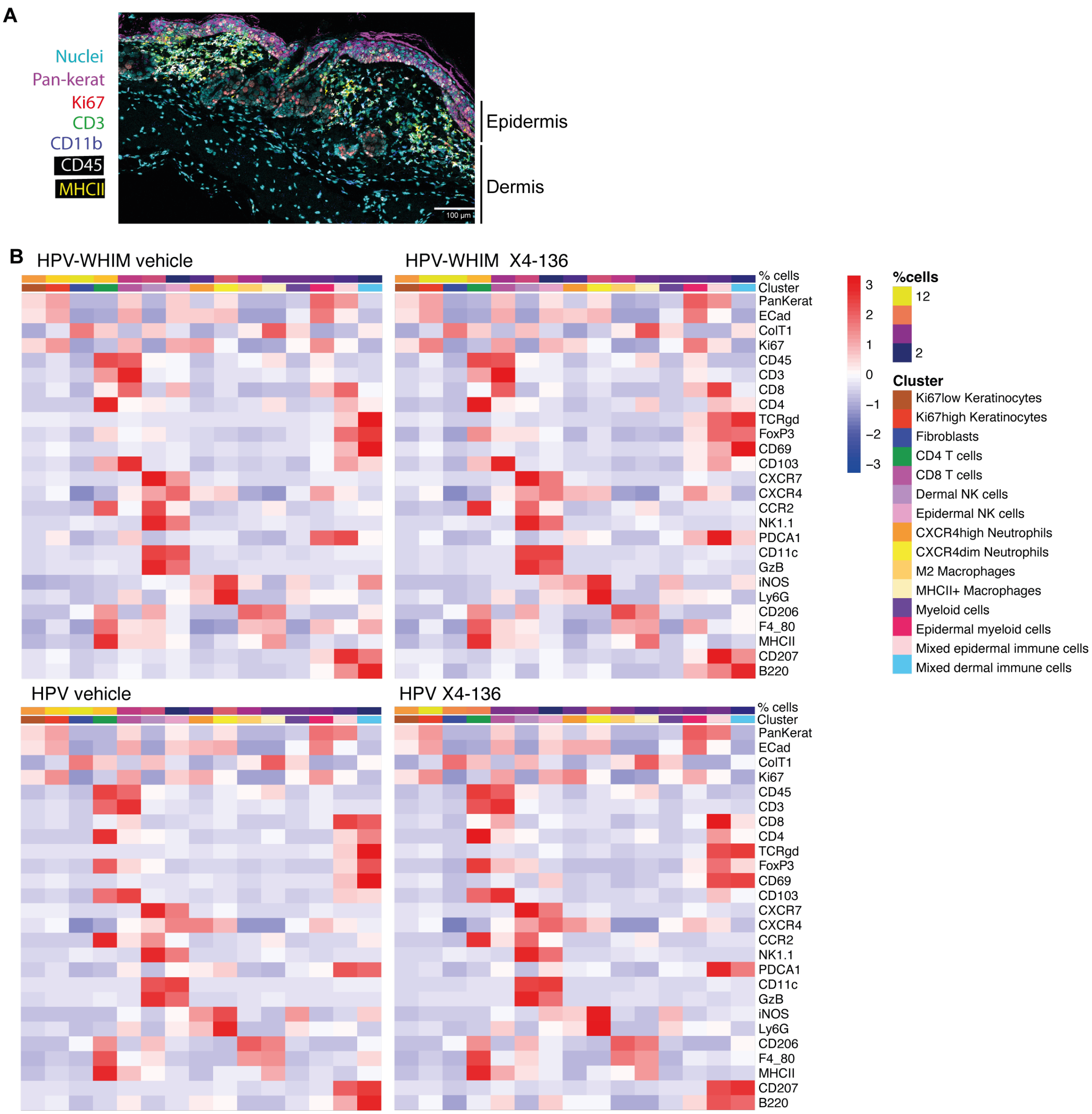


**Supplemental Figure 4. Unsupervised IMC analysis identifies 15 clusters. (A)** Representative image of a region of interest (ROI) defined for IMC analysis. Six markers and the nuclei are shown. Epidermis and dermis are indicated. Scale bar = 100 µm. **(B)** Heatmap visualization of the mean expression of population, activation, and proliferation markers for each cell type for all four experimental groups. The proportions (%) of cells in each cluster are shown at the top of each heatmap. **(A-B)** Data from 2 experiments, with a total of 5 mice /group.

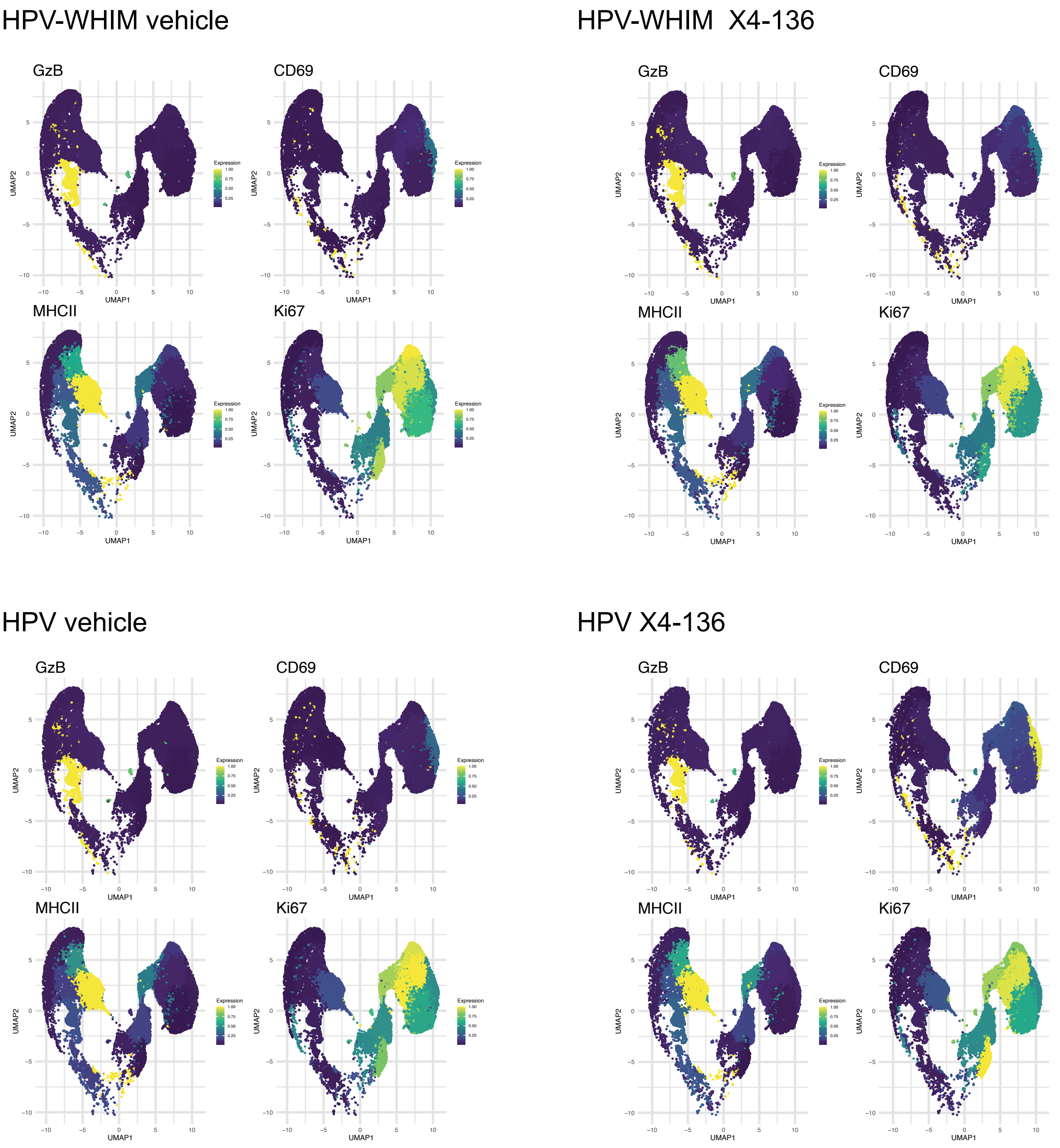


**Supplemental Figure 5. CXCR4 inhibition with the X4-136 compound maintains the overall phenotypic profile of cell subtypes.** UMAP visualization of single cells from the skin of mice treated or not by X4-136, colored by the mean marker expression (GzB, MHCII, CD69, Ki767) of each cell type. A statistical comparison of the expression of activation and proliferation markers does not show any significant differences between groups. Data from 2 experiments, with a total of 5 mice /group. Statistical analysis was performed using the Kruskal-Wallis test, with Benjamini-Hochberg p-value correction to account for multiple testing in the unsupervised analysis. *$p < 0.05$.

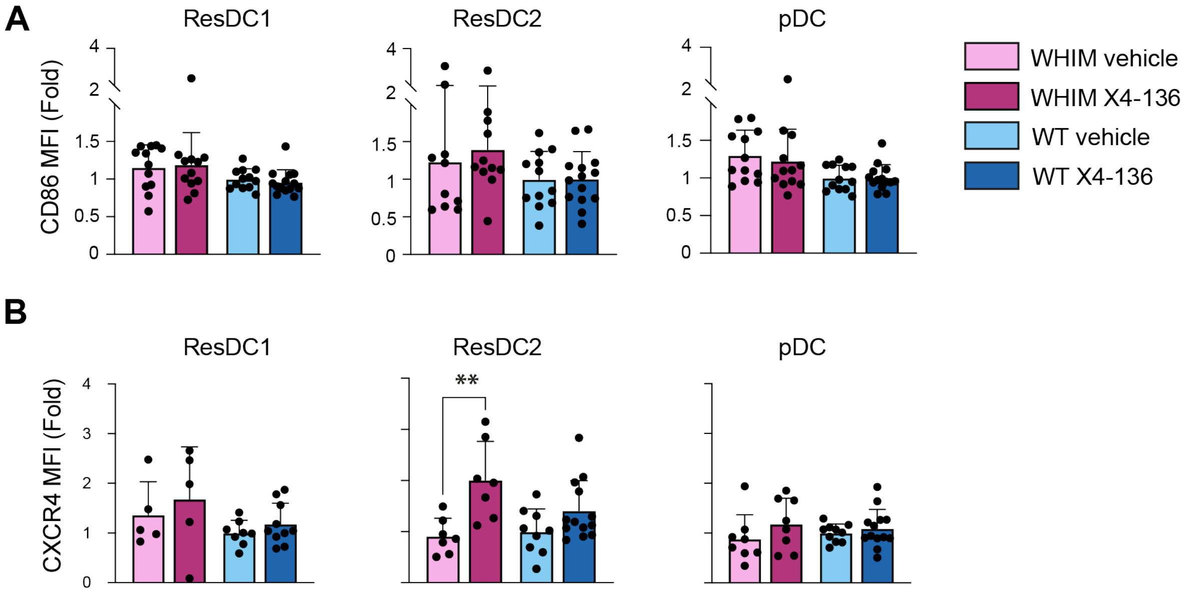


**Supplemental Figure 6. Impact of CXCR4 inhibition with the X4-136 compound on the phenotype of resDCs and pDCs recovered in SDLNs in acute inflammation. (A)** CD86 and **(B)** CXCR4 membrane expression levels in total resDC1 (left panels), resDC2 (middle panel), and pDCs (right panel) from vehicle- and X4-136-treated WT and WHIM mice. Data are expressed as the fold change compared to the mean MFI in each experiment's "WT vehicle" group. Bar graphs show the mean ± SD for cumulative data from 4 to 5 experiments, with a total of 5-14 mice/group. Statistical analysis was performed using the Kruskal-Wallis test. **$p < 0.01$.

**Supplementary Table 1**

| Antigen | Fluorochrome | Supplier | Clone | Reference | Dilution |
|---|---|---|---|---|---|
| Viability dye | eFluor 506 | eBiosciences | NA | 65-0866-14 | 1/1000 |
| CD3 | Alexa Fluor 700 | BD Biosciences | 17A2 | 561388 | 1/100 |
| CD3e | eFluor 450 | eBiosciences | 145-2C11 | 48-0031-82 | 1/100 |
| CD4 | APC-eFluor 780 | eBiosciences | GK1.5 | 47-0041-82 | 1/100 |
| CD8α | APC-eFluor 780 | eBiosciences | 53-6.7 | 47-0081-82 | 1/100 |
| CD8α | BV605 | BD Biosciences | 53-6.7 | 563152 | 1/100 |
| CD11b | PerCP Cy5.5 | BD Biosciences | M1/70 | 550993 | 1/200 |
| CD11b | AlexaFluor 700 | BD Biosciences | M1/70 | 557960 | 1/200 |
| CD11c | APC | eBiosciences | N418 | 17-0114-81 | 1/100 |
| CD11c | PE-Cy7 | eBiosciences | N418 | 25-0114-82 | 1/100 |
| CD16/CD32 | Purified | BD Biosciences | 2.4G2 | 553142 | 1/50 |
| CD19 | PE-CF594 | BD Biosciences | 1D3 | 562291 | 1/250 |
| CD24 | APC-eFluor 780 | eBiosciences | M1/69 | 47-0242-82 | 1/100 |
| CD45 | FITC | eBiosciences | 30-F11 | 11-0451-82 | 1/100 |
| CD64 | PE-Dazzle™ 594 | BioLegend | X54-5/7.1 | 139320 | 1/200 |
| CD86 | BV605 | BD Biosciences | GL1 | 563055 | 1/250 |
| CD86 | PE | BD Biosciences | GL1 | 553692 | 1/250 |
| CD103 | APC-R700 | BD Biosciences | M290 | 565529 | 1/100 |
| CD103 | PE | BD Biosciences | M290 | 557495 | 1/100 |
| CD206 | APC | BD Biosciences | MR5D3 | 565250 | 1/100 |
| CXCR4 | APC | eBiosciences | 2B11 | 17-9991-82 | 1/100 |
| Rat IgG2bκ | APC | eBiosciences | EB149/10H5 | 17-4031-81 | 1/100 |
| F4/80 | BV605 | BD Biosciences | T45-2342 | 743281 | 1/100 |
| Ly6C | FITC | BD Biosciences | AL-21 | 553104 | 1/100 |
| Ly6G | PE-Cy7 | BD Biosciences | 1A8 | 560601 | 1/250 |
| MHC class II | PerCP-Vio700 | Miltenyi Biotec | M5/114.15.2 | 130-103-805 | 1/200 |
| PDCA1 | BV421 | BioLegend | 927 | 127023 | 1/100 |

**Supplementary Table 2**

| Antigen | Tag | Supplier | Clone | Reference | Dilution |
|---|---|---|---|---|---|
| ACKR3 | 167Er | Proteintech | 4C3D7 | 60216-1-Ig | 1/400 |
| B220 | 147Sm | BioLegend | RA36B2 | 103202 | 1/200 |
| CCR2 | 156Gd | Abcam | EPR20844-15 | ab273050 | 1/100 |
| CD103 | 152Sm | Abcam | EPR22590-27 | ab224202 | 1/50 |
| CD11b | 149Sm | Standard Biotools | EPR1344 | 3149028D | 1/600 |
| CD11c | 162Dy | Standard Biotools | N418 | 3162017B | 1/25 |
| CD19 | 142Nd | Standard Biotools | 6OMP31 | 3142014D | 1/200 |
| CD206 | 164Dy | Abcam | EPR25215-277 | ab300621 | 1/3000 |
| CD207 | 171Yb | Abcam | EPR24685-12 | ab283686 | 1/50 |
| CD3 | 170Er | Standard Biotools | Polyclonal | 3170019D | 1/200 |
| CD4 | 169Tm | Standard Biotools | BLR167J | 91H031169 | 1/100 |
| CD45 | 151Eu | Standard Biotools | D3F8Q | 91H029151 | 1/400 |
| CD69 | 145Nd | Standard Biotools | H1.2F3 | 3145005B | 1/50 |
| CD8 | 176Yb | Standard Biotools | EPR21769 | 91H023176 | 1/200 |
| Collagen type I | 141Pr | Standard Biotools | Polyclonal | 91H018141 | 1/750 |
| CXCR4 | 163Dy | eBioscience | 2B11 | 14-9991-82 | 1/50 |
| E-Cadherin | 174Yb | Standard Biotools | 24E10 | 91H011174 | 1/300 |
| F4/80 | 153Eu | Standard Biotools | D2S9R | 91H030153 | 1/75 |
| FoxP3 | 158Gd | Standard Biotools | FJK-16s | 91H032158 | 1/100 |
| Granzyme B | 155Gd | Standard Biotools | EPR22645-206 | 91H026155 | 1/600 |
| iNOS | 160Gd | Standard Biotools | SP126 | 91H025160 | 1/200 |
| Ki67 | 173Yb | Standard Biotools | B56 | 91H017173 | 1/400 |
| Ly-6G | 166Er | Standard Biotools | 1A8 | 91H037166 | 1/500 |
| MHC class II | 161Dy | Standard Biotools | M5/114.15.2 | 91H038161 | 1/100 |
| NK1.1 | 165Ho | Standard Biotools | PK136 | 3165018B | 1/250 |
| Pan-cytokeratin | 148Nd | Standard Biotools | AE-1/AE-3 | 3148022D | 1/1000 |
| PDCA1 | 175Lu | Novus Bio | 120G8.04 | DDX0390P-100 | 1/200 |
| TCRß | 143Nd | Standard Biotools | H57-597 | 3143010B | 1/50 |
| TCRγδ | 159Tb | Standard Biotools | GL3 | 3159012B | 1/25 |
